# *Drosophila centrocortin* is dispensable for centriole duplication but contributes to centrosome separation

**DOI:** 10.1101/2021.08.31.458397

**Authors:** Dipen S. Mehta, Pearl V. Ryder, Jina Lee, Hala Zein-Sabatto, Dorothy A. Lerit

**Author notes:** These authors contributed equally (Dipen S. Mehta, Pearl V. Ryder).

## Abstract

Centrosomes are microtubule-organizing centers that duplicate exactly once to organize the bipolar mitotic spindle required for error-free mitosis. Prior work indicated that *Drosophila centrocortin* (*cen*) is required for normal centrosome separation, although a role in centriole duplication was not closely examined. Through time-lapse recordings of rapid syncytial divisions, we monitored centriole duplication and the kinetics of centrosome separation in control versus *cen* null embryos. Our data suggest that although *cen* is dispensable for centriole duplication, it contributes to centrosome separation.

## Description

Centrosomes are the microtubule-organizing centers (MTOCs) of most eukaryotic cells that promote error-free mitosis through organization of the bipolar mitotic spindle. Centrosome function as a MTOC is instructed by the pericentriolar material (PCM), a matrix of proteins that encircles the central pair of centrioles (Nigg and Raff 2009). RNAs also localize to centrosomes, and local mRNAs may contribute to centrosome functions, likely through a co-translational transport mechanism (Marshall and Rosenbaum 2000; Sepulveda *et al.* 2018; Chouaib *et al.* 2020; Kwon *et al.* 2021; Safieddine *et al.* 2021). For example, recent work indicates *centrocortin (cen)* mRNA localized to centrosomes is required for error-free mitosis in rapidly dividing, syncytial *Drosophila* embryos (Bergalet *et al.* 2020; Ryder *et al.* 2020). Historically, several hypotheses have been suggested to account for why mRNAs reside at centrosomes, including the postulation that mRNA may help instruct centrosome duplication (Alliegro *et al.* 2006). Normally, centrioles duplicate only once per cell cycle, and this regulation is key to prevent multipolar spindles and chromosomal instability (Wong and Stearns 2003; Tsou and Stearns 2006).

*cen* mutant embryos null for protein and mRNA show mitotic defects, including errant centrosome separation and multipolar spindles, despite normal microtubule assembly (Kao and Megraw 2009). However, centriole duplication was not examined in *cen* mutants. To address whether *cen* contributes to centriole duplication, we examined the kinetics of centrosome duplication and separation in live control versus *cen* null *Drosophila* embryos co-expressing GFP-Cnn to label the PCM and RFP-PACT to mark the centrioles (**Figure 1A**). Through blinded image analysis, we calculated the time elapsed from when centrosomes are duplicated, visible as two proximal RFP-PACT signals, until they were fully separated (**Figure 1B**). Centrosomes from *cen* null embryos duplicated and initiated centriole separation (N=45 centrosomes from N=3 mutant embryos). However, about 9% of *cen* centrosomes (4/45 centrosome pairs; open dots, **Figure 1B**) failed to fully separate, consistent with prior work (Kao and Megraw 2009). In contrast, all observed centrioles duplicated and separated normally in control embryos (N=32 centrosomes from N=2 embryos). Although the rate of centrosome separation, as shown in **Figure 1B**, was not significantly different in *cen* mutants relative to controls (time to separate, expressed as mean + S.D., was 344.4 + 83.8 s for *cen* mutants versus 306.3 + 65.8 s for controls), a chi-square test did indicate that the centrosomes in *cen* mutant embryos require more time (>400 s) to separate (**Figure 1C**; * *p*<0.05). These data together with the failure of some centrosomes to complete centrosome separation suggests *cen* is dispensable for centriole duplication but contributes to normal centrosome separation. Further work is required to unearth precisely how *cen* promotes centrosome separation.

**Figure 1.**
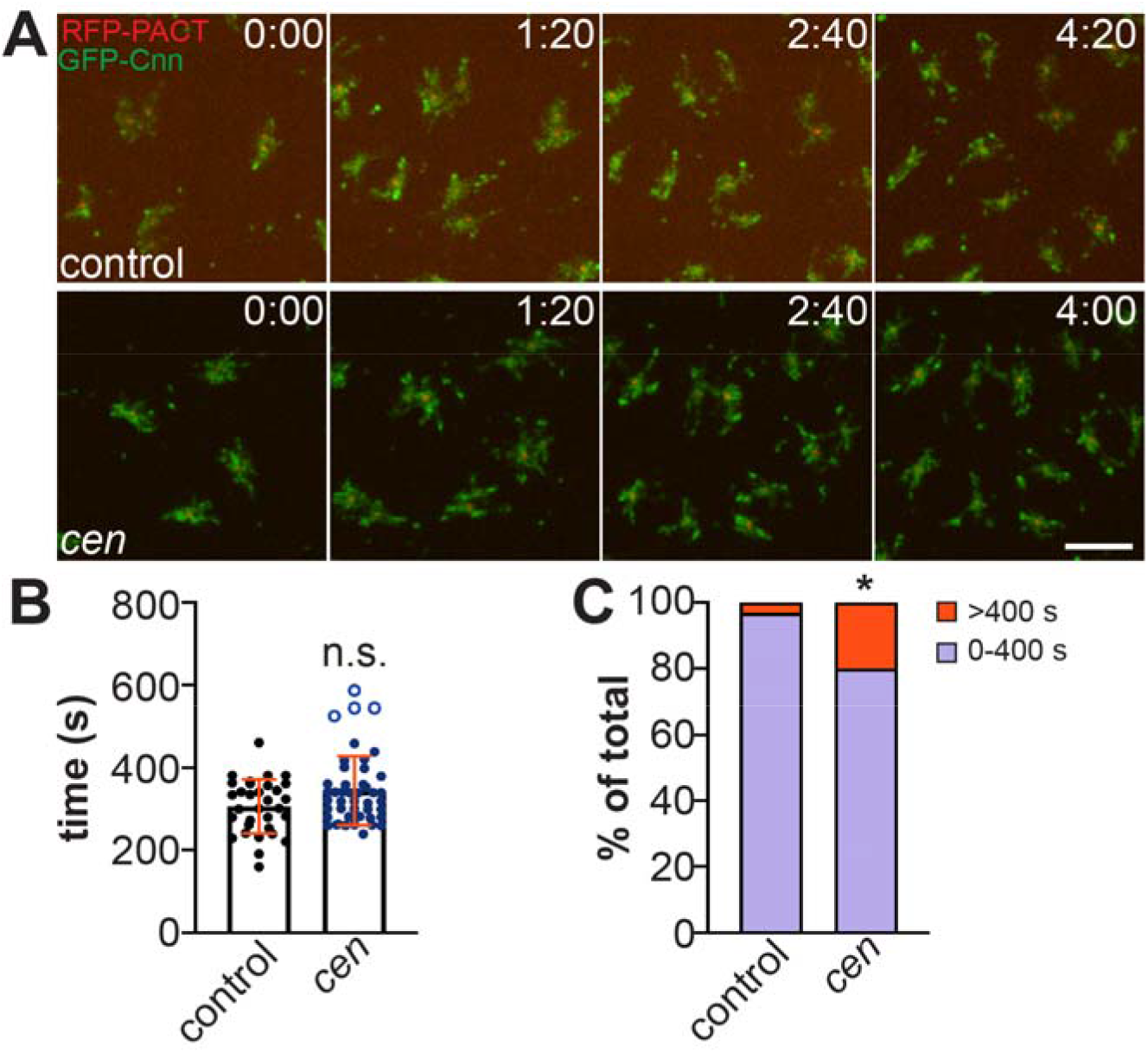
*cen* promotes centrosome separation. (A) Stills from time-lapse recordings of NC 12 control and *cen* mutant embryos expressing RFP-PACT and GFP-Cnn. Time is shown in min:s. (B) Quantification shows the time (s) required to fully separate the duplicated centrosomes from control versus *cen* mutant NC 12 embryos. Each dot represents a single measurement from WT: N=32 centrosomes from 2 embryos and *cen*: N=45 centrosomes from 3 embryos. Open dots represent centrosomes that never completed separation prior to the end of the recording. n.s., not significant by a two-tailed Mann-Whitney test, *p*=0.171. (C) Distribution analysis of the time to complete centrosome separation, as detailed in (B). A chi-square test indicates that centrosomes from *cen* mutant embryos take significantly longer time (>400 s) to separate relative to controls, *X*^2^ (1, N = 77) = 4.7, \**p*=0.030. Bar, 5 μm.

## Methods

*P{Ubi-RFP-PACT}* was a gift from Nasser Rusan (NIH) and was generated by introducing amino acids 2,479–3,555 from PLP-PF (Fragment 5), into the vector *pURW* (stock 1282, *Drosophila Genomics Resource Center*) to generate an RFP fusion as described for the GFP fusions in (Galletta *et al.* 2014). Transgenic animals were generated by BestGene, Inc. (Chino Hills, CA). Flies were raised on Bloomington formula ‘Fly Food B’ (LabExpress; Ann Arbor, MI), and crosses were maintained at 25°C in a light and temperature-controlled incubator chamber. To examine maternal effects, mutant embryos are progeny derived from mutant mothers.

For live imaging, dechorionated 1-2 hour embryos were adhered to a 22×30 mm #1.5 coverslip using glue extracted from double-sided tape (3M), covered with Halocarbon oil 700 (H8898; Millipore-Sigma), and inverted onto a 50-mm gas permeable dish (Sarstedt) with broken #1 coverslips used as spacers. Images were captured on a Nikon Ti-E inverted microscope fitted with Yokagawa CSU-X1 spinning disk head using a motorized stage, Nikon LU-N4 solid state laser launch (15 mW 405, 488, 561, and 647 nm), Hamamatsu Orca Flash 4.0 v2 digital CMOS camera, and a Nikon 100x, 1.49 NA Apo TIRF oil immersion objective. Images were acquired at 300 ms exposure times over 0.5 mm Z-intervals through 3.5 mm of tissue at 20 s time intervals for one or more complete embryonic nuclear cycles.

Genotype-blinded image analysis was completed in FIJI (NIH; (Schindelin *et al.* 2012)). Data were plotted and statistical analysis was performed using Microsoft Excel and GraphPad Prism software.

## Reagents

| *Drosophila* strain | Source | Description |
| --- | --- | --- |
| *P<sub>BAC</sub>-GFP-Cnn* | Lerit et al. (2015) | Cnn tagged at the N-terminus with EGFP under endogenous regulatory elements |
| *P{Ubi-RFP- PACT}* | This paper | N-terminal fusion of RFP to a C-terminal fragment of PLP containing the PACT domain |
| *cen*<sup>f04787</sup> | Kao and Megraw (2009) | Protein and RNA null allele; Bloomington stock #18805 |

## Acknowledgments

We thank Nasser Rusan for gifts of fly strains and Beverly Robinson for technical advice. Stocks obtained from the Bloomington *Drosophila* Stock Center (NIH grant P40OD018537) and reagents from the *Drosophila* Genomics Resource Center (NIH grant 2P40OD010949) were used in this study.

## Author Contributions

Dipen S. Mehta: formal analysis, writing - review editing. Pearl V. Ryder: investigation, writing - review editing, conceptualization. Jina Lee: investigation. Hala Zein-Sabatto: investigation, formal analysis. Dorothy A. Lerit: conceptualization, funding acquisition, resources, supervision, visualization, writing - original draft, writing - review editing.

## References

Alliegro MC, Alliegro MA, Palazzo RE. 2006. Centrosome-associated RNA in surf clam oocytes. Proc Natl Acad Sci U S A 103: 9034–8. PubMed

Bergalet J, Patel D, Legendre F, Lapointe C, Benoit Bouvrette LP, Chin A, Blanchette M, Kwon E, Lécuyer E. 2020. Inter- dependent Centrosomal Co-localization of the cen and ik2 cis-Natural Antisense mRNAs in Drosophila. Cell Rep 30: 3339–3352.e6. PubMed

Chouaib R, Safieddine A, Pichon X, Imbert A, Kwon OS, Samacoits A, Traboulsi AM, Robert MC, Tsanov N, Coleno E, Poser I, Zimmer C, Hyman A, Le Hir H, Zibara K, Peter M, Mueller F, Walter T, Bertrand E. 2020. A Dual Protein-mRNA Localization Screen Reveals Compartmentalized Translation and Widespread Co-translational RNA Targeting. Dev Cell 54: 773–791.e5. PubMed

Galletta BJ, Guillen RX, Fagerstrom CJ, Brownlee CW, Lerit DA, Megraw TL, Rogers GC, Rusan NM. 2014. Drosophila pericentrin requires interaction with calmodulin for its function at centrosomes and neuronal basal bodies but not at sperm basal bodies. Mol Biol Cell 25: 2682–94. PubMed

Kao LR, Megraw TL. 2009. Centrocortin cooperates with centrosomin to organize Drosophila embryonic cleavage furrows. Curr Biol 19: 937–42. PubMed

Kwon OS, Mishra R, Safieddine A, Coleno E, Alasseur Q, Faucourt M, Barbosa I, Bertrand E, Spassky N, Le Hir H. 2021. Exon junction complex dependent mRNA localization is linked to centrosome organization during ciliogenesis. Nat Commun 12: 1351. PubMed

Lerit DA, Jordan HA, Poulton JS, Fagerstrom CJ, Galletta BJ, Peifer M, Rusan NM. 2015. Interphase centrosome organization by the PLP-Cnn scaffold is required for centrosome function. J Cell Biol 210: 79–97. PubMed

Marshall WF, Rosenbaum JL. 2000. Are there nucleic acids in the centrosome? Curr Top Dev Biol 49: 187–205. PubMed

Nigg EA, Raff JW. 2009. Centrioles, centrosomes, and cilia in health and disease. Cell 139: 663–78. PubMed

Ryder PV, Fang J, Lerit DA. 2020. centrocortin RNA localization to centrosomes is regulated by FMRP and facilitates error-free mitosis. J Cell Biol 219: e202004101. PubMed

Safieddine A, Coleno E, Salloum S, Imbert A, Traboulsi AM, Kwon OS, Lionneton F, Georget V, Robert MC, Gostan T, Lecellier CH, Chouaib R, Pichon X, Le Hir H, Zibara K, Mueller F, Walter T, Peter M, Bertrand E. 2021. A choreography of centrosomal mRNAs reveals a conserved localization mechanism involving active polysome transport. Nat Commun 12: 1352. PubMed

Schindelin J, Arganda-Carreras I, Frise E, Kaynig V, Longair M, Pietzsch T, Preibisch S, Rueden C, Saalfeld S, Schmid B, Tinevez JY, White DJ, Hartenstein V, Eliceiri K, Tomancak P, Cardona A. 2012. Fiji: an open-source platform for biological-image analysis. Nat Methods 9: 676–82. PubMed

Sepulveda G, Antkowiak M, Brust-Mascher I, Mahe K, Ou T, Castro NM, Christensen LN, Cheung L, Jiang X, Yoon D, Huang B, Jao LE. 2018. Co-translational protein targeting facilitates centrosomal recruitment of PCNT during centrosome maturation in vertebrates. Elife 7: e34959. PubMed

Tsou MF, Stearns T. 2006. Mechanism limiting centrosome duplication to once per cell cycle. Nature 442: 947–51. PubMed

Wong C, Stearns T. 2003. Centrosome number is controlled by a centrosome-intrinsic block to reduplication. Nat Cell Biol 5: 539–44. PubMed

